# A single-cell atlas of breast cancer cell lines to study tumour heterogeneity and drug response

**DOI:** 10.1101/2021.03.02.433590

**Authors:** G Gambardella, G Viscido, B Tumaini, A Isacchi, R Bosotti, D di Bernardo

## Abstract

Breast cancer patient stratification is mainly driven by tumour receptor status and histological grading and subtyping, with about twenty percent of patients for which absence of any actionable biomarkers results in no clear therapeutic intervention. Cancer cells within the same tumour have heterogeneous phenotypes and exhibit dynamic plasticity. However, how to evaluate such heterogeneity and its impact on outcome and drug response is still unclear. Here, we transcriptionally profiled 35,276 individual cells from 32 breast cancer cell lines covering all main breast cancer subtypes to yield a breast cancer cell line atlas. We found high degree of heterogeneity in the expression of clinically relevant biomarkers across individual cells within the same cell line; such heterogeneity is non-genetic and dynamic. We computationally mapped single cell transcriptional profiles of patients’ tumour biopsies to the atlas to determine their composition in terms of cell lines. Each tumour was found to be heterogenous and composed of multiple cell lines mostly, but not exclusively, of the same subtype. We then trained an algorithm on the atlas to determine cell line composition from bulk gene expression profiles of tumour biopsies, thus providing a novel approach to patient stratification. Finally, we linked results from large-scale in vitro drug screening^1,2^ to the single cell data to computationally predict responses to more than 450 anticancer agents starting from single-cell transcriptional profiles. We thus found that transcriptional heterogeneity enables cells with differential drug sensitivity to co-exist in the same population. Our work provides a unique resource and a novel framework to determine tumour heterogeneity and drug response in breast cancer patients.

## Introduction

One of the main roadblocks to personalized medicine of cancer is the lack of biomarkers to predict outcome and drug sensitivity from a tumour biopsy. Multigene assays such as MammaPrint^3^, Oncotype DX^4,5^ and PAM50^6^ can classify Breast Cancer (BC) tumour types and risk of relapse^7^ but with limited clinical utility^7,8^. Genomic and transcriptional biomarkers of drug sensitivity are available only for a restricted number of drugs^1,2,9^. As a consequence, BC patient stratification is still mainly driven by receptor status and histological grading and subtyping^7^, with about twenty percent^10^ of patients for which paucity of actionable biomarkers limits personalized therapies. Moreover, even when a targeted treatment option is available, drug resistance may arise^7^ partly because of rare drug tolerant cells characterized by distinct transcriptional or mutational states^11–17^.

Determining tumour heterogeneity and its impact on drug response is essential to better stratify patients and aid in the development of personalized therapies. Expression-based biomarkers measured from bulk RNA-sequencing of a tumour biopsy are powerful predictors of drug response in vitro^1,2,18^, but average out tumour heterogeneity. Single-cell transcriptomics yields a molecular profile of each cell^19,20^, however, it is still unclear if and how it can inform clinical decision making. Here, we focused on tumour-derived breast cancer cell lines. We hypothesized that despite being simplistic models of tumours, cancer cell lines may exhibit themselves heterogeneous phenotypes, and serve as cell-state “primitives” to deconvolve tumour cell composition from patients’ biopsies for patient stratification and prediction of drug response.

## RESULTS

### 1. Single-cell Transcriptome Profiling of Breast cancer cell lines

We performed single cell RNA-sequencing (scRNA-seq) of 31 breast cancer cell lines (Supplementary Table 01) and one non-cancer cell line, MCF12A^21^, by means of the Drop-seq technology^20^. Following pre-processing (Methods), we retained a total of 35,276 cells, with an average of 1,069 cells per cell line and 3,248 genes captured per cell (Supplementary Figure 01 and Supplementary Table 01).

We next generated an atlas (http://bcatlas.tigem.it) encompassing the 32 BC cell lines, as shown in Figure 1A. In the atlas, luminal BC cell lines form a big “island” with multiple “peninsulas” with intermixing of cells from distinct cell lines; on the contrary, triple-negative breast cancer (TNBC) cell lines give rise to an “archipelago”, where cells tend to separate into distinct islands according to the cell line of origin, thus suggesting that TNBC cell lines represent instances of distinct diseases.

**Figure 1.**
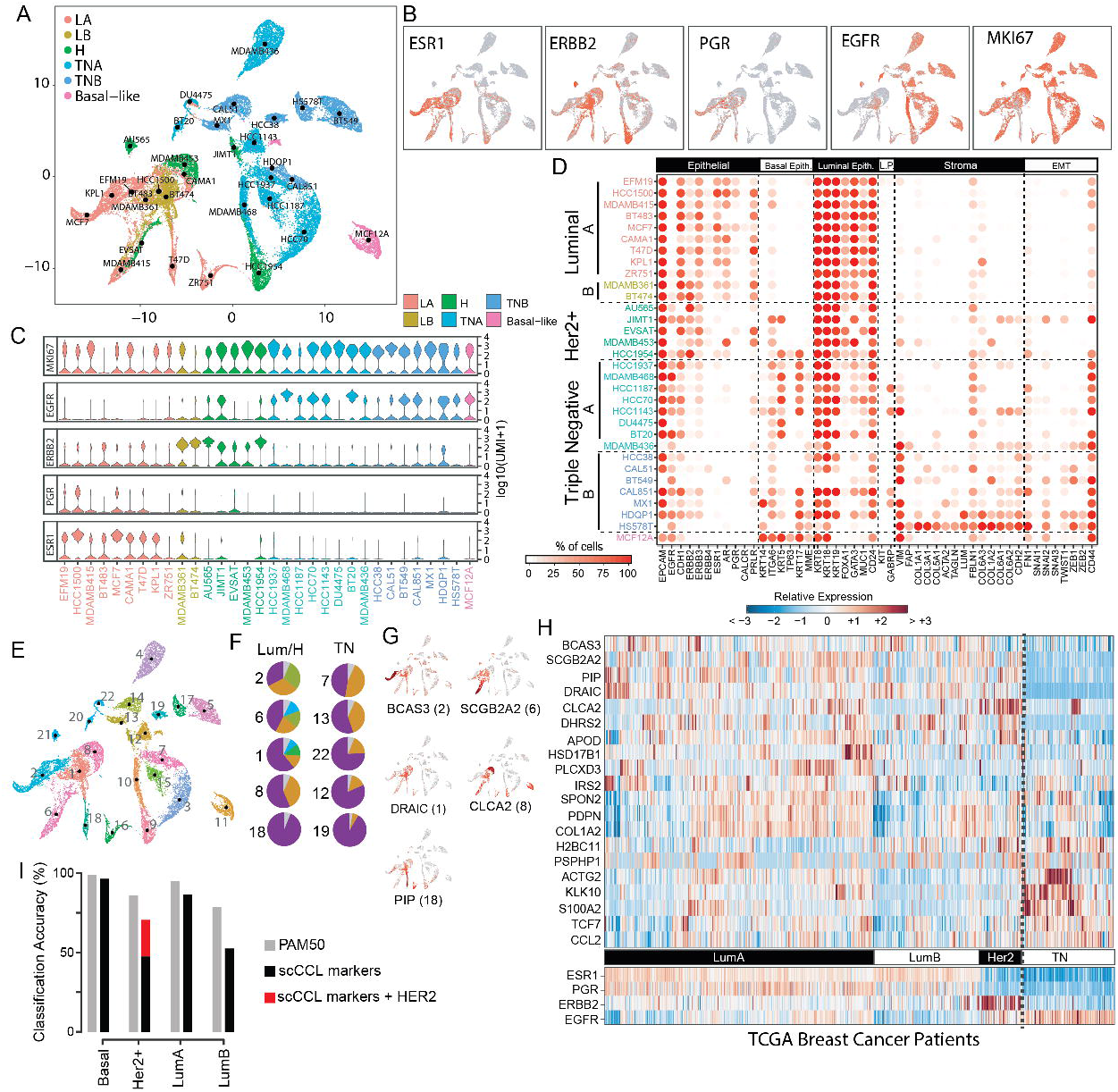
The Breast Cancer Single Cell Atlas. (**A**) Representation of single-cell expression profiles of 35,276 cells from 32 cell lines color-coded according to cancer subtype (LA=Luminal A, LB=Luminal B, H=Her2 positive, TNA = Triple Negative A, TNB = Triple Negative B). (**B**) Expression levels of the indicated genes in the atlas, with red indicating expression, together with their (**C**) distribution within the cell lines, shown as a violin plot. (**D**) Dotplot of literature-based biomarker genes along the columns for each of the 32 sequenced cell lines along the rows. Biomarker genes are grouped by type (Basal Epith. = Basal Epithelial, Luminal Epith. = Luminal Epithelial, L.P. = Luminal Progenitor, EMT = Epithelial to Mesenchymal Transition). (**E**) Graphical representation of 35,276 cells color-coded according to their cluster of origin. Clusters are numbered from 1 to 22. (**F**) For the indicated cluster, the corresponding pie-chart represents the cluster composition in terms of cell lines. Cell lines in the same pie-chart are distinguished by colour. Only the top 10 most heterogenous clusters are shown. Cluster 2 is the most heterogeneous while cluster 19 is the most homogeneous. (**G**) Expression levels in the atlas of the five luminal biomarkers identified as the most differentially expressed in each of the five luminal clusters (1, 2, 6, 8 and 18). (**H**) Expression of 20 out of 22 atlas-derived biomarkers in the biopsies of 937 breast cancer patient from TCGA. (**I**) Accuracy in classifying tumour subtype for 937 patients from TCGA by using either PAM50 or the 20 atlas derived biomarker genes (scCCL) alone or augmented with HER2 gene (scCCL + HER2).

Single-cell expression of clinically relevant biomarkers (Figure 1B,C) including oestrogen receptor 1 (ESR1), progesterone receptor (PGR), Erb-B2 Receptor Tyrosine Kinase 2 (ERBB2 a.k.a. HER2) and the epithelial growth factor receptor (EGFR) across the different cell lines are in agreement with their reported status^21–23^.

To gain further insights into each cancer cell line, we analysed the expression of 48 literature-based biomarkers of clinical relevance^24^, as reported in Figure 1D. Luminal cell lines highly express luminal epithelium genes, but neither basal epithelial nor stromal markers; on the contrary, triple-negative BC cell lines (11 out of 15) show a basal-like phenotype with the expression of at least one of keratin 5, 14 or 17^25,26^, with triple-negative subtype B (TNB) cell lines also expressing vimentin (VIM) and Collagen Type VI Alpha Chains (COL6A1, COL6A2, COL6A3)^21^. Interestingly, two out of five HER2 overexpressing (HER2^+^) cell lines (JIMT1 and HCC1954) in the atlas are in the triple-negative “archipelago” and express keratin 5 (KRT5) (Figure 1A,D), which has been linked to poor prognosis and trastuzumab resistance^27^. Indeed, both cell lines are resistant to anti-HER2 treatments^28^. Finally, the non-tumorigenic MCF12A cell line lacks expression of ESR1, PGR and HER2 and displays a basal-like phenotype characterized by the expression of all basal-like marker genes including keratin 5, 14, 17 and TP63, in agreement with the literature^29^.

Overall, these results show that single cell transcriptomics can be successfully used to capture the overall expression of clinically relevant markers.

### 2. The BC single-cell atlas identifies clinically relevant transcriptional signatures

By clustering the 35,276 single-cells in the atlas, we identified 22 clusters, as shown in Figure 1E. Within the luminal island, cells did not cluster according to their cell line of origin, indeed four out of the five luminal clusters contain cells from distinct cell lines (Figure 1F and Supplementary Figure 02). On the contrary, triple-negative cell lines clustered according to their cell line of origin, with each cluster containing mostly cells from the same cell line (Figure 1F).

We identified genes specifically expressed among cells in the same cluster for a total of 22 biomarkers, one for each cluster (Figure 1G,H and Supplementary Figure 03). Interestingly, neither *ESR1* nor *ERRB2* were part of this set. Literature mining confirmed the significance of some of these markers: clusters in the luminal island (Figure 1G) were associated to genes involved in cancer progression (BCAS3^30,31^ cluster 2), dissemination (SCGB2A2^32,33^ cluster 6), proliferation (DRAIC^34,35^ cluster 1), migration and invasion (CLCA2^36,37^ cluster 8 and PIP^38^ cluster 18). Interestingly, whereas DRAIC is correlated with poorer survival of luminal BC patients^35^, both CLCA2 and PIP are significantly associated with a favourable prognosis^36,37,39,40^.

To examine the clinical relevance of these 22 biomarkers, we analysed their expression across 937 breast cancer patients from the TGCA collection encompassing all four BC types. Out of the 22 biomarkers, two (MAGEA4 and XAGE2) could not be mapped to the TGCA dataset. As shown in Figure 1H, there is a marked difference in the expression of the 20 cluster-derived biomarkers across Luminal A, Luminal B, Her2 positive and Triple Negative patients. Moreover, it is possible to distinguish subtypes within each category, which may lead to novel diagnostic/prognostic biomarkers (Figure 1H and Supplementary Figure 04). For example, one subset of triple-negative patients strongly expresses the protease kallikrein-10 (KLK10), which has been associated with poor prognosis, poor response to tamoxifen treatment^41^ and identified as potential target to reverse trastuzumab resistance^42^. Whereas a second subset is characterised by actin gamma 2 expression (ACTG2), which has been linked in BC to cell proliferation^43^ and platinum-based chemotherapy sensitivity^44–47^.

Finally, we compared the performance of the 20 biomarker genes in classifying BC subtypes from bulk RNA-seq data (Methods) against the PAM50 gene signature (50 genes)^6^ used in clinics to identify breast cancer subtypes (Figure 1I). The performances were overall comparable, with the obvious exceptions of HER2-overexpressing cancers. Indeed, when adding *ERBB2* to the list of 20 cluster-based biomarkers, classification of this subtypes markedly improved (Figure 1I).

Altogether, these analyses confirm that the single cell BC cell line atlas allows identifying clinically relevant gene signatures useful for patient stratification and tumour type classification.

### 3. The BC atlas as a reference for automated cancer diagnosis

The BC atlas can be used as a reference against which to compare single cell transcriptomics data from a patient’s tissue biopsy and to perform cancer subtype classification and assessment of tumour heterogeneity. To this end, we developed an algorithm able to map single-cell transcriptional profiles from a patient onto the BC atlas and to assign a specific cell line to each of the patient’s cells (Methods). We first tested the ability of the algorithm in correctly classifying the very cells in the atlas starting from their single-cell transcriptional profiles and correctly classified 92% of the cells (Supplementary Figure 05). We then turned to single-cell transcriptional profiles obtained from five triple-negative breast cancer patients^48^. As shown in Figure 2A, most, but not all the patients’ cells mapped to the triple-negative “archipelago”, except for the TNBC5 sample, for which most cells mapped to the luminal island. As the algorithm assigns a specific cell line to each tumour cell, it is also possible to look at the cell line composition of each patient, as reported in Figure 2B. These results demonstrates that heterogeneity varies across patients but is present in all the samples, as no patient’s biopsy mapped to a single cell line. Moreover, information on the drug sensitivity of the individual cell lines composing the tumour may prove useful in guiding therapeutic choices.

**Figure 2.**
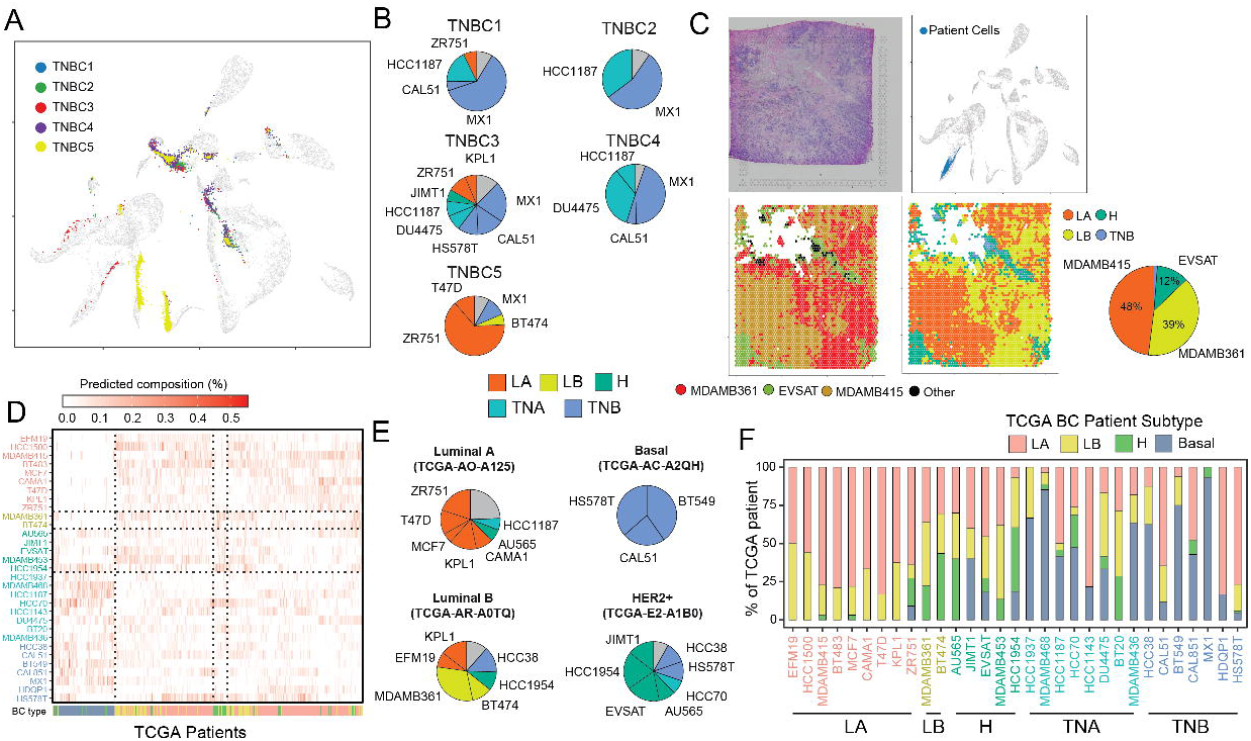
Automatic classification of patients’ tumour cells. (**A**) Cancer cells from triple negative breast cancer (TNBC) biopsies of 5 patients are embedded in the BC atlas to predict their tumour type. (**B**) For each patient, the pie chart shows cell line composition obtained by mapping patient’s cells onto the atlas. (**C**) Tissue-slide of an oestrogen positive breast tumour biopsy sequenced using 10x Visium spatial transcriptomics (top-left) and the position of the mapped tissue tiles onto the atlas (top-left). Tiles are colour-coded according to the cell line (bottom-left) and to tumor subtype (bottom-right) as predicted by the mapping algorithm. (**D**) Cell line composition for each patient as estimated by the algorithm from bulk RNA-seq of 937 BC patients. For ease of interpretation, in the heatmap patients are clustered according to their cell line composition. The bottom row reports the annotated cancer subtype in TGCA. (**E**) Predicted cell-line composition for four representative patients. (**F**) The distribution of the 937 BC patients across the 32 cell lines. For each cell line, the stacked bars report the percentage of patients of a given cancer subtype assigned by the algorithm to that cell line.

We next tested the algorithm on spatial transcriptomics dataset obtained from the tissue biopsy of two patients, one diagnosed with ESR1^+^/ERBB2^+^ lobular oestrogen positive carcinoma (Figure 2C-E and Supplementary Figure 06A) and the other with ESR1^+^/ERBB2^+^ ductal carcinoma (Supplementary Figure 06C,D)^49^. The dataset consists of 3,808 transcriptional profiles for patient 1 (Figure 2C) and 3,615 profiles for patient 2 (Supplementary Figure 06C), each obtained from a different tissue “tile” of size 100um x 100um x 100 um. The algorithm projected each of the spatial tiles onto the BC atlas and assigned a cell line to each tile. We coloured the tiles according to the cell line and the BC subtype of the cell line (Figure 2C) to yield an automatic cancer subtype classification of tiles. Most of the tiles for both patients were assigned to just two cell lines and correctly classified as luminal (A or B); the remaining 13% of the tiles for patient 1 and 20% for patient 2 were instead classified either as HER2-overexpressing or Triple Negative, which could be an important information to guide therapeutic choice and to predict the occurrence of drug resistance.

As bulk gene expression profiles are more clinically relevant than single-cell gene expression profiles, we next trained a deconvolution algorithm <u>Bisque</u>^50^ (Methods and Supplementary Figure 07) by leveraging our single-cell atlas to predict the cell line composition of a tumour sample. To test the effectiveness of this algorithm, we collected 937 bulk gene expression profiles from breast cancer patients in TGCA whose BC subtypes were annotated, and then assigned to each patient the corresponding cell line composition, as shown in Figure 2D,E. Reassuringly, patients diagnosed with a specific breast cancer subtype tend to have a tumour cell line composition consisting of cell lines of the same subtype. We quantified this observation in Figure 2F and observed some interesting exceptions: JIMT-1 is an HER2-overexpressing cell line with an amplified ERBB2 locus, but no HER2+ patient was mapped to this cell line. Interestingly, JIMT-1 cells are resistant to anti-HER2 treatments^51^; another example is the HS578T cell line, which is reported to be triple-negative, however the majority of patients who map to it are luminal; surprisingly, this cell line has been reported to be sensitive to fulvestrant^1,2^, an anti-ESR1 drug.

These results show that this single cell atlas of cancer cell can be used to automatically assign cell line composition and cancer subtypes both from single-cell expression profiles and bulk gene expression profile.

### 4. Clinically relevant biomarkers exhibit heterogenous and dynamic expression in BC cell lines

Clinically relevant receptors are heterogeneously expressed across cells belonging to the same cell line, as assessed by computing the percentage of cells in a cell line expressing the receptor as in Figure 3A. Consider the seven Luminal B and HER2^+^ cell lines present in the BC atlas, which by definition overexpress HER2: whereas more than 90% of cells in AU565, BT574 and HCC1954 cell lines express *ERBB2*, in the remaining four cell lines *ERBB2* expression ranged from 31% of EVSAT cells to 46% of JIMT1 cells and up to 64% of MDA-MB-361 cells. This happens despite both JIMT1 and MDA-MB-361 harbour a copy number gain of the locus containing the *ERBB2*^52^. We first excluded the possibility that these results were artifacts of single-cell RNA-sequencing technology (Supplementary Figure 08). We then assessed HER2 protein levels by flow cytometry in three representative cell lines: AU565 (high HER2 expression), MDA-MB-361 (heterogeneous HER2 expression) and HCC38 cell lines (low HER2 expression). As shown in Figure 3B, single-cell transcriptional data agree with the cytometric analysis; however, the origin of this heterogeneity is unclear. To exclude hereditable genetic differences as a source of heterogeneity, we sorted MDA-MB-361 cells into HER2^+^ and HER2^-^ subpopulations (Methods) and checked whether these homogenous subpopulations were stable over time, or rather spontaneously gave rise to heterogeneous populations. As shown in Figure 3C, after 18 days in culture, both subpopulations re-established the original heterogeneity, demonstrating that HER2 expression in these cells is dynamic and driven by a yet undiscovered mechanism.

**Figure 3.**
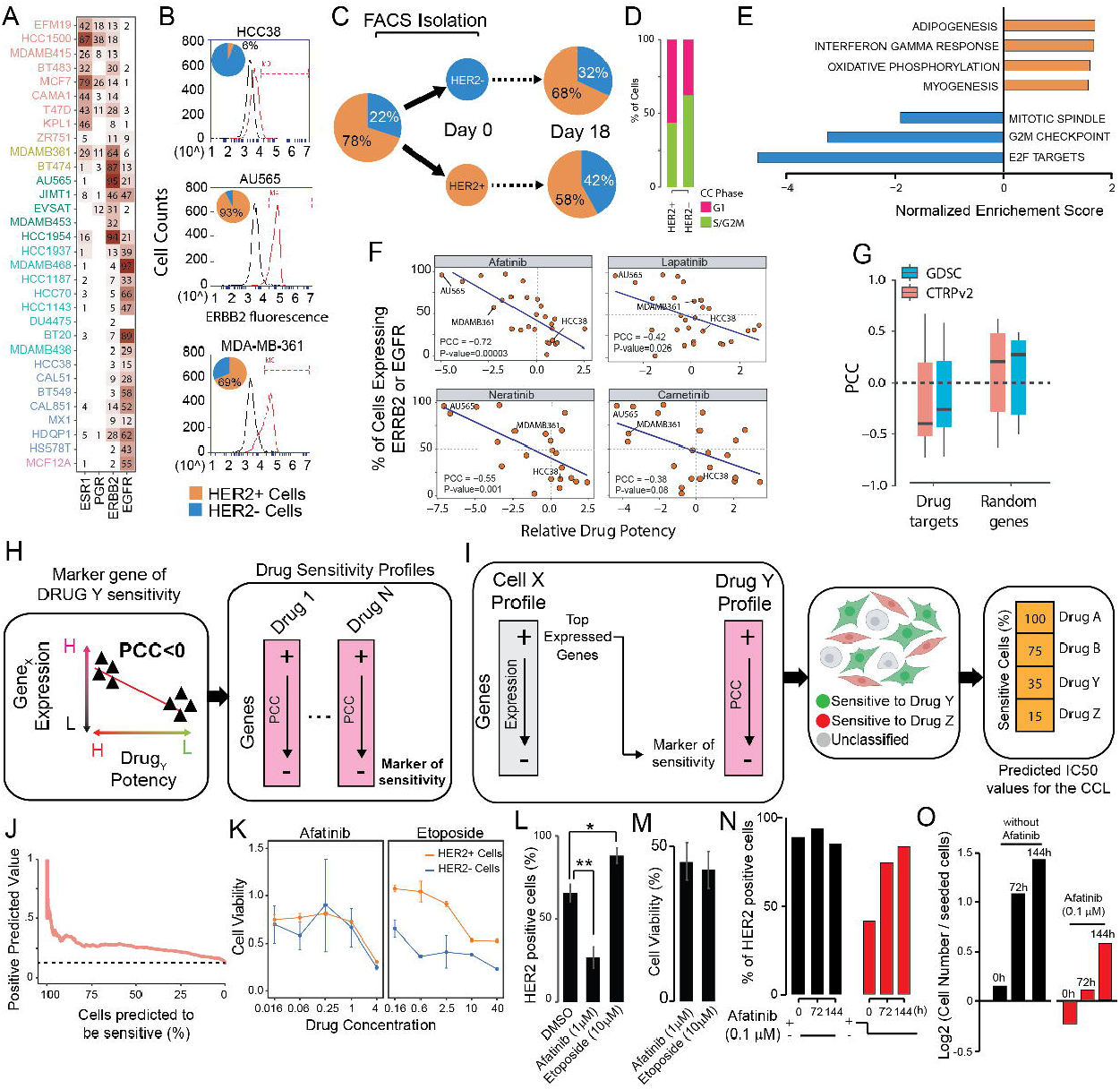
Transcriptional heterogeneity in breast cancer cell lines and its impact on drug response. (**A**) Percentage of cells expressing the indicated genes in each of the sequenced 32 cell lines. (**B**) Fluorescence cytometry of HCC38, MDA-MB-361 and AU565 cell lines stained with a fluorescent antibody against Her2. (**C**) Expression of HER2 protein in MDA-MB-361 cells is dynamic and re-established in less than 3 weeks. (**D**) Analysis of the cell cycle phase for the HER2+ and HER2-subpopulations of MDA-MB-361 cells. The cell cycle of each cell is estimated from its single-cell transcriptomics profile. (**E**) Enriched pathways (GSEA, FDR<10%) across the genes differentially expressed between the HER2+ and HER2-subpopulations of MDA-MB-361 cells. Orange refers to HER2+ subpopulation and blue to the HER2-ones. (**F**) Relationship between gene expression and drug potency for four anti-HER2 drugs. Each dot corresponds to a cell line reporting the percentage of cells expressing ERBB2 or EGFR in the cell line [y-axis] and the drug potency [x-axis]. PCC (pearson correlation coefficient) and p-value are also shown. (**G**) Box-plot reporting the distribution of PCCs between percentage of cells expressing the cognate drug target and the potency of the drug across cell lines for 66 drugs for two different drug potency databases. For comparison, the PCC distribution when choosing a random gene in place of the cognate drug target is also shown. (**H**) Bioinformatics pipeline for the identification of drug sensitivity biomarkers for 450 drugs. For each drug, the expression of a gene across 658 cell lines is correlated with drug potency in the same cell lines; genes are then ranked from most positively correlated to the most negatively correlated. (**I**) The top 250 most expressed genes in a single cell are used as input for a Gene Set Enrichment Analysis (GSEA) against the ranked list of genes for each one of the 450 drugs to predict its drug sensitivity. At the end of the process, each cell in the sample is associated to the drug it is most sensitive to, or to no drug, if no significant enrichment score is found. Finally, for each of the 450 drugs, the number of cells predicted to be either sensitive, resistant, or not classified in the considered sample is estimated. (**J**) Validation of DREEP on the Breast Cancer Single Cell atlas data to predict drug sensitivity to 86 drugs. The PPV (Positive Predicted Value) is shown as a function of the percentage of cells in a cell line predicted to be sensitive to the same drug. Dashed line represents the performance of a random algorithm. (**K**) Dose-response curve for afatinib and etoposide on sorted MDA-MB-361 cell populations (triplicate experiment). (**L**) Percentage of HER2+ cells in MDA-MB-361 after 72h treatment with either afatinib (statistic: two-sided t-test, *P ≤ 0.05; **P ≤ 0.01; ***P ≤ 0.001) or etoposide and (**M**) measured cell viability after the treatment. (**N**) Percentage of HER2 positive cells in MDA-MB-361 cell-line at the indicated time-points either after 48h of afatinib pre-treatment (red bars) or without any afatinib pre-treatment (black bars) and (**O**) the relative number of cells rescaled for the number of cells at the beginning of the experiment.

Interestingly, HER2^+^ circulating tumour cells (CTCs) isolated from an ER^+^/HER2^-^ breast cancer patient were shown to spontaneously interconvert from HER2^-^ and HER2^+^, with cells harbouring a phenotype producing daughters of the opposite one^53^. To check if cell-cycle phase could explain the observed heterogeneity in the MDA-MB-361 cell line, we computationally predicted (Methods) the cell cycle phase of each cell in both the HER2^-^ and HER2^+^ subpopulations from single cell transcriptomics data^54^. A higher proportion of HER2^-^ cells was predicted to be in S/G2/M phases when compared to HER2+ cells (Figure 3D). This result is consistent with previous observations that report cell cycle arrest in G2/M phase following HER2 inhibition^55^.

We next set to identify biological processes differing between the two subpopulations by computing differentially expressed genes (DEGs) from the single-cell transcriptional profiles of HER2^+^ cells against HER2^-^ cells (Supplementary Table 02). Gene Set Enrichment Analyses (GSEA) ^56^ against the ranked list of DEGs, reported in Figure 3E, revealed seven significantly enriched pathways (FDR<10%): four of which were upregulated in HER2^+^ cells, but downregulated in HER2^-^ cells, and included adipogenesis, myogenesis and OXPHOS, all indicative of EMT engagement, which has been reported in HER2^+^ cells^57–59^; the remaining three pathways were upregulated in HER2^-^ cells and related to cell-cycle and specifically to G2/M phase, in agreement with our previous analysis, suggesting that cell cycle may play a role in HER2 expression in this cell line.

These results show that heterogeneity in the expression of clinically relevant biomarkers is present even in cell lines and that it can also be dynamic and of a non-genetic nature.

### 5. Heterogeneity in gene expression affects drug response

To investigate the role of heterogeneity in gene expression within a cell line on drug response, we collected large-scale in vitro drug screening data^1,2^ reporting the effect of 450 drugs on 658 cancer cell lines from solid tumours. As show in Figure 3F and Supplementary Figure 09, sensitivity of the BC cell lines to HER2 inhibitors was significantly correlated with the percentage of cells in the cell line expressing *ERBB2* (Supplementary Table 03). Receptor expression level is substantially the same across cells expressing it, irrespective of the cell line they belong to (Supplementary Figure 10), except for cell lines harbouring CNVs of the *ERBB2* locus. Furthermore, we found that the correlation between drug target expression and drug sensitivity holds true also for several other targets (Figure 3G), thus suggesting that variability in gene expression within cells of the same tumour may cause some cells to respond poorly to the drug treatment.

Starting from these observations, we developed DREEP (DRug Estimation from single-cell Expression Profiles), a novel bioinformatics tool that, starting from single-cell transcriptional profiles, allows to predict drug response at the single cell level. To this end, we first detected expression-based biomarkers of drug sensitivity for 450 drugs^2^, as schematised in Figure 4H,I (Methods). Briefly, we crossed data from the Cancer Cell Line Encyclopaedia (CCLE) on the response to 450 drugs across 658 cancer cell lines from solid tumours with their gene expression profiles from bulk RNA-seq. In the CCLE, drug potency is evaluated as the inverse of the Area Under the Curve (AUC) of the dose-response graph, with low values of the AUC indicating drug sensitivity, while high values implying drug resistance (Figure 3H). For each gene and for each drug, we computed the correlation between the expression of the gene across the 658 cell lines with the drug potency in the same cell lines. Hence, genes positively correlated with the AUC are potential markers of resistance, vice-versa, negatively correlated genes are markers of sensitivity (Figure 3H). In this way, we generated a ranked list of expression-based biomarkers of drug sensitivity and resistance for each of the 450 drugs. We then used these biomarkers to predict drug sensitivity at the single-cell level (Figure 3I). To this end, the 250 genes most expressed of each cell in the atlas were compared against the ranked list of biomarkers for each one of 450 drugs by means of GSEA^56^ and thus associated to the drug it is most sensitive to, or to no drug, if no significant enrichment score from GSEA is found (Figure 3I).

To assess the algorithm’s performance, we applied it to the single-cell BC atlas and estimated its performance by checking how well we could predict sensitivity of the 32 BC cell lines to 86 drugs for which this information was publicly available^60^ (Figure 3J). To convert single-cell predictions to predictions at the cell line level, we simply used the percentage of cells in the cell line deemed to be sensitive to the drug by the algorithm. To experimentally validate DREEP, we turned to the MDA-MB-361 cell line for which we found coexistence of two distinct and dynamic cell subpopulations (HER2^+^ and HER2^-^). We applied DREEP to each subpopulation to identify drugs able to selectively inhibit growth of either the HER2^-^ subpopulation or the HER2^+^ subpopulation: 42 drugs (FDR < 1%, Supplementary Table 04) were predicted to preferentially inhibit growth of HER2^-^ cells; the most overrepresented class among these drugs was that of inhibitors of DNA topoisomerases (TOP1/TOP2A) (Supplementary Figure 11) such as Etoposide. Surprisingly, no drug was found to specifically inhibit growth of HER^+^ cells, whereas 44 drugs (FDR <1%) were predicted to be equally effective on both subpopulations and unexpectedly included HER2 inhibitors, such as afatinib (Supplementary Table 03 and Supplementary Figure 12).

We selected etoposide and afatinib for further experimental validation. MDA-MB-361 cells were first sorted by FACS into HER2^+^ and HER2^-^ subpopulations and then cell viability was measured following 72h drug treatment at five different concentrations as shown in Figure 3K (and Supplementary Table 05). In agreement with DREEP predictions, HER2^-^ cells were much more sensitive to etoposide than HER2^+^ cells, while afatinib was equally effective on both subpopulations. This counterintuitive result was similar to that observed by Jordan et al^53^ using circulating tumour cells from a BC patient sorted into HER2^-^ and HER2+ subpopulations, which were found to be equally sensitive to Lapatinib (another HER2 inhibitor), but no mechanism of action was put forward.

We hypothesise that the dynamic interconversion of MDA-MB-361 cells between the HER2^-^ and the HER2^+^ state may explain this surprising result: when the starting population consists of HER2^-^ cells only, some of these cells will nevertheless interconvert to HER2^+^ cells during afatinib treatment, and they will thus become sensitive to HER2 inhibition, explaining the observed results. We mathematically formalised this hypothesis with a simple mathematical model (Supplementary Figure 13 and in the Supplementary Material) where two species (HER2^+^ and HER2^-^ cells) can replicate and interconvert, but only one (HER2^+^) is affected by afatinib treatment. The model shows that if the interconversion time between the two cell states is comparable to that of the cell cycle, then afatinib treatment will have the same effect on both subpopulations. If instead the interconversion time is much longer than the cell cycle, then afatinib will have little effect on HER2^-^ sorted cells, but maximal effects on HER2^+^ sorted cells, and vice-versa, if the interconversion time is much shorter than the cell cycle, then afatinib’s effect would be minimal on both HER2^-^ and HER2^+^ sorted cells.

Comparison of the modelling results with the experimental results thus suggests that the interconversion rate should be of the same order of the cell cycle (about 72h for MDAM361 cells). The model further predicts that treating the unsorted population of MDA-MB-361 cells with afatinib reduces the percentage of HER2^+^ cells, since only HER2^+^ will be affected, but that this percentage quickly recovers once Afatinib treatment is interrupted (Supplementary Figure 14 and 15 and Supplementary Material).

To test modelling predictions, we treated the MDAM361 cell line (without sorting) with afatinib and etoposide and then assessed by cytofluorimetry the percentage of HER2+ and HER2^-^ cells before and after the treatment. As shown in Figure 3L,M (Supplementary Table 06 and Supplementary Table 07) etoposide increased the percentage of HER2^+^ cells, in agreement with the increased sensitivity of HER2^-^ cells to this treatment, whereas afatinib strongly decreased the percentage of HER2^+^ cells, confirming that its effect is specific for HER2^+^ cells only. We next measured the percentage of HER2^+^ cells following removal of afatinib from the medium; as shown in Figure 3N,O the percentage of HER2+ cells quickly increased confirming the modelling results (Supplementary Figure 15 and Supplementary Material).

All together our results show that DREEP can predict drug sensitivity from single-cell transcriptional profiles and that dynamic heterogeneity in gene expression does play a significant role in how the cell population will respond to the drug treatment.

## Discussion

In this study we provide the first transcriptional characterization at single cell level of a panel of 32 breast cell lines. We show that single cell transcriptomics can be used to capture the expression of clinically relevant markers. We show that breast-cancer cell lines express clinically relevant BC receptors heterogeneously among cells within the same cell line. Moreover, we observed dynamic plasticity in the regulation of HER2 expression in the MDA-MB-361 cell line with striking consequences on drug response. This phenomenon has been recently observed also in circulating tumour cells of a BC patient^53^ and in other cell lines^17,61^.

We determined cell line composition of patients’ biopsies both from both single-cell and bulk gene expression profiles. Estimation of cancer cell line composition provides an alternative and more information-rich framework to link bulk gene expression measurement of patient’s biopsies to preclinical cancer models. Knowledge of drugs to which cancer cell lines are sensitive to may also inform drug treatment for patients for which bulk gene expression profiles have been measured.

Single cell transcriptomics is still not clinically ready because of the costs and time needed, however this work shows the importance of performing single-cell sequencing on the available cancer models, including cell lines and organoids to build a set of cell cancer states with known phenotypes and drug response to which patients’ tumour can be mapped to make a leap in personalised diagnosis, prognosis and treatment of cancer patients.

## Methods

### Cell culture

The 32 cell lines used in this study were obtained from commercial providers and cultured in ATCC recommended complete media at 37°C and 5% CO2.

### DROP-seq platform set-up

Single cell transcriptomic of the 32 cell lines was performed by implementing in-house the DROP-seq technology^20^. The microfluidics device for the generation of droplet was fabricated using a bio-compatible, silicon-based polymer, polydimethylsiloxane (PDMS) that was rendered hydrophobic with Aquapel® treatment as per protocol^20^. In each sequencing experiment, cell suspension, bead suspension and carrier oil (QX200 droplet generation oil, Bio-Rad) were first loaded in syringes and then placed in syringe pumps (Leafluid). Flow rates of syringe pumps were set at 4,000 μL/hr for both cell and barcoded bead suspensions while carrier oil syringe pump was set at 15,000 μL/hr. In each sequencing experiment, cells and barcoded beads were respectively diluted at the concentration of 200 cell/μL in PBS with BSA 0.01% (Merck) and 120 bead/μL in lysis buffer. A self-built magnetic stirrer system was used to keep in suspension barcoded beads. To count the occurrence of a single cell together with a barcoded bead several tests were performed without lyses buffer in the bead suspension. In these tests, we observed about 5% of generated droplets filled with just one bead and one cell.

### Single cell RNA library preparation and sequencing

For each sequencing experiment, the targeted number of cells to sequence was set to 2,000. Droplets were collected in a 50 mL falcon and broke by adding 1 mL of Perfluoro-1-octanol. Captured RNA was reverse transcribed in a single reaction following the original protocol ^20^ and then digested with exonuclease 1 to degrade unbound primers. Next, cDNA was first amplified with a total of 12 PCR cycles and then purified using AMPure XP beads at 0.6X ratio. Finally, the quality of the resulting cDNA library was quantified with the BioAnalyzer High Sensitivity DNA Chip and its concentration measured using the Qubit Fluorometer. The Illumina Nextera XT v2 kit was used to produce the next generation sequencing (NGS) libraries using four aliquots of 600pg of each cDNA library. Quality and concentration of NGS libraries were respectively quantified on the BioAnalyzer High Sensitivity DNA Chip and Qubit Fluorometer. Finally, either Illumina NextSeq 500/550 or NovaSeq 6000 machines were used to sequence the produced NGS libraries (Supplementary Table 01). Samples processed with NextSeq500/550 NGS library were diluted at the final concentration of 3 nM and sequenced using the 75-cycle high output flow cell while samples processed with NovaSeq 6000 machine were diluted at the final concentration of 250 pM and sequenced using the S1 100 cycles flow cell.

### Read alignment and gene expression quantification

Raw data processing was performed using the Drop-seq tools package version 1.13 and following the Drop-seq Core Computational Protocol (http://mccarrolllab.org/dropseq). Briefly, raw sequence data was filtered to remove all read pairs with at least one base in their barcode or UMI with a quality score less than 10. Then read 2 was trimmed at the 5’ end to remove any TSO adapter sequence, and at the 3’ end to remove polyA tails. Reads were then aligned using STAR ^62^ on hg38 human genome (primary assembly, version 28) downloaded from GENCODE ^63^. After reads alignment, UMI tool ^64^ was used to perform UMI deduplication and quantify the number of gene transcripts in each cell. The initial number of sequenced cells was identified using a simple (knee-like) filtering rule as implemented by CellRanger 2.2.x. After this, only high depth cells with at least 2,500 UMI, more than 1,000 captured genes and with less than 50% of reads aligned on mitochondrial gene were retained. Putative multiples among the sequenced cells of each BC cell line were simply discarded identifying outliers in the count depth distribution by using Tukey’s method based on lower and upper quartiles with k equal to 3.

### BC Atlas Construction

Single cells expression profiles were normalized using GF-ICF (Gene Frequency – Inverse Cell Frequency) normalization using the *gficf* package^65,66^ for R statistical environment (https://github.com/dibbelab/gficf). GF-ICF is based on a data transformation model called term frequency-inverse document frequency (TF-IDF) that has been extensively used in the field of text mining. GF-ICF transformation was applied on CPM (count per million) after *EdgeR* normalization ^67^ and discarding genes expressed in less than 5% of the total number of sequenced cells. Finally, each cell was summarized with its first 10 Principal Components (PCs) and projected with UMAP ^68^ into a two dimensional embedded space. The number of principal components was chosen as the “elbow” point on the plot of the first 50 PCs. UMAP projection was performed by using the *uwot* package in the R statistical environment 3.6.

### Cell clustering and identification of marker genes

Transcriptionally similar subpopulations of cells were found using a Phenograph like approach^69^ as implemented in the *clustcells* function of *gficf* package^65^. Briefly, we initially built a graph of cells by using the K-Nearest Neighbours (KNN) algorithm applied to the PC-reduced space where each cell was connected to its 50 most similar cells using the manhattan distance. Then, to build the final graph of cells, the edge weight between any two cells was computed as the Jaccard similarity, i.e. the proportion of neighbours they share. The Louvain algorithm with resolution parameter equal to 0.25 was used to find communities of cells in this graph. Differentially expressed genes in each cluster were identified by the *findClusterMarkers* function of *gficf* package, which compares the expression of a gene in each cluster versus all the other by using the Wilcoxon rank-sum test^65^.

### TGCA bulk expression dataset and cell-line deconvolution

Raw bulk expression data and relative patient clinical information were collected from the Genomic Data Commons (GDC) portal^70^ by using the *TCGAbiolinks* package^71^. Then, raw counts were normalized using the *EdgeR* package^67^ into R statistical environment 3.6. Bisque tool^50^ (available at https://github.com/cozygene/bisque) was used to estimate the cell-line composition from the patient’s bulk gene expression profile. Specifically, we applied the *ReferenceBasedDecomposition* function with parameters: *bulk.eset* set to the bulk gene expression dataset in log2 scale; *sc.eset* set to our single-cell BC atlas with normalized raw counts rescaled in log2; *use.overlap* set to FALSE and *markers* set to the marker genes across the 32 BC cell-lines estimated by using the function *findClusterMarkers* of *gficf* package. As in the original manuscript describing the Bisque tool^50^, only marker genes with an FDR<0.5 and Log2 fold change greaten then 0.25 were used for deconvolution purpose.

### Spatial sequencing data

Spatial transcriptomic data of two BC patients were download from 10x Genomic website (https://www.10xgenomics.com/resources/datasets). Only tiles reported to be “in tissue” according to the related metadata of each patient slide were used.

### Mapping new cells into the BC atlas and estimation of the cancer subtype

New points were mapped to the UMAP space via *embedNewCells* function of *gficf* package^65^. Briefly, tiles from 10x spatial transcriptomics were normalized with *gficf* package using the ICF weight estimated on the BC atlas. Then tiles were projected to the existing PC space using gene loadings from the BC atlas. After this transformation, tiles were mapped to the BC atlas via *umap_transform* function of *uwot* package. Finally the cancer subtype of each mapped tile was predicted with the function *classify.cells* of the package *gficf* with the k nearest-neighbour parameter set to 7.

### Single-cell drug sensitivity prediction

The naïve gene expression profile (RNA-seq) of about 1,000 cancer cell line was obtained from the Cancer Cell Line Encyclopaedia (CCLE) portal^72^. Cell lines belonging to liquid tumour were discarded and only 658 cell lines belonging to solid tumours were retained and used for further analysis. The raw counts of each gene were normalized with edgeR package ^67^ and transformed in log10(CPM+1). Poorly expressed genes and genes whose entropy was in the fifth percentile were excluded from the analysis. Expression profiles of the 658 CCLs were then crossed with drug sensitivity data^2^. This dataset was originally composed of 481 small molecules, but, after removing drugs for which the in vitro response was available for less than 25 CCLs, only 450 small molecules were retained for further analysis. For each gene and for each of the 450 drugs, we computed the Pearson correlation coefficient (PCC) between the expression of the gene across the 658 cell lines and the effect of the drug expressed in terms of Area Under the Curve (AUC). Since the AUC reflects the in vitro response of a cell line to different concertation of a drug in a timeframe of 72 hours, lower values of AUC are associated with sensitivity whereas higher values with resistance to the drug. Hence, genes positively correlated with the AUC are potential markers of resistance (the more expressed the gene, the higher the concentration needed to inhibit growth), vice-versa, negatively correlated genes are markers of sensitivity. We this approach, we generated a ranked list of expression-based biomarkers of drug sensitivity and resistance for each of the 450 drugs where genes positively correlated with the AUC are at the top, and those negatively correlated at the bottom. Finally, to predict drug sensitivity at the single-cell level, we used the top 250 expressed genes of each cell as input of Gene Set Enrichment Analysis (GSEA) ^56^ against the ranked list of biomarkers for each one of 450 drugs built as described above. Hence, while a negative enrichment score implies that genes associated to drug sensitivity are highly expressed by the cell, a positive one indicates the cell express genes conferring drug resistance. GSEA and associated p-values were estimating using the *fgsea* package in the R statistical environment version 3.6.

### Drug sensitivity of the HER2+ and HER2-subpopulations in the MDA-MB-361 cell line

For each sequenced cell of the MDA-MB-361 cell line, the enrichment score of 450 anticancer drugs was predicted as described above. Then, to identify drugs exhibiting differential sensitivity for the two subpopulations, we used the Mann-Whitney test was to assess if there was a difference between the enrichment scores of HER2+ and HER2-subpopulations. P-values were corrected for false discovery rate using Benjamini-Hochberg correction. A drug was considered specific for HER2-cell population if and only if its FDR was less than 0.05 and the median enrichment score across HER2-cells less than zero while its median enrichment score across HER2+ cells greater than zero. Conversely, a drug was considered specific for HER2+ cell population if and only if FDR was less than 0.05 and the median enrichment score across HER2+ cells less than zero while its median enrichment score across HER2-cells greater than zero.

### Validation of drug sensitivity prediction

Precision of the DREEP method in predicting drug sensitivity from single cell transcriptional profiles was evaluated using an independent publicly available drug screening dataset^9^ composed by 1,001 CCLs and their maximal inhibitory concentration (IC50) values for 265 small molecules. Hence, we applied DREEP to the single-cell profiles of the 32 BC cell lines to predict the percentage of sensitive cells in each cell line for the 86 drugs. The “golden standard” was built by assigning to each of 32 × 86 (=2,752) cell line/drug pair the value 1 if the cell line was sensitive to the drug and 0 otherwise. To determine if a cell line was sensitive or not to a specific drug from the experimental data, we converted for each drug its IC50 distribution in Z-scores using all the 1,001 available cell lines and then defined a cell line sensitive to the drug if and only if its Z-score was in the 5% percentile. Finally, Positive Predicted Values (PPV) were defined as TP/(TP+FP) where TP represents the number of true positives and FP the number of false positives predicted cell lines/drug pairs.

### Prediction of cell cycle phase from scRNA-seq

The cell cycle phase of each sequenced cell was predicted using the function *CellCycleScoring* of the *Seurat* tool with default parameter and following what was suggested in the corresponding vignette (https://satijalab.org/seurat).

### HER2 antibody staining procedure for flow cytometry analysis

Cells were first washed with phosphate-buffered saline (PBS) 1x, detached with 0.05% trypsin-EDTA, resuspended and harvested with the appropriate medium in single-cell suspension. Then, cells were counted, washed with PBS-FBS 1%, and finally incubated for 15 min at 4° in the dark at the concentration of 1.0 × 10^6^ cell/μL with staining buffer. The staining buffer was prepared diluting the mouse anti-human HER2 antibody (BD BB700) at the final concentration of 0.00114 ng/μL. Then, to remove unbound antibody, cells were washed three times with PBS-FBS 1%. Flow cytometry measurements were performed on either BD Accuri C6 or BD FACSAria III instruments. To define antibody positive and negative cells, the unstained samples were used to set the gate. To record data, at least 1.0 × 10^4^ events were collected for each sample. Data analysis was performed using the either BD FACSDiva 8.0.1 or BD Accuri C6 software.

### HER2 expression dynamics experiment

Sorting of MDA-MB-361 HER2-positive and HER2-negative cells was performed following the antibody staining procedure described above with the only exception that before sorting, each sample was resuspended in sorting buffer (PBS 1x, FBS 1%, trypsin 0.1%, EDTA 2mM). Then, 4.0 × 10^5^ cells were collected for each cell subpopulation (*i.e*. HER2-positive and HER2-negative), plated in their appropriate medium, and incubated at 37°. After 18 days, the percentage of cells expressing HER2 protein was checked by performing the antibody staining procedure described above.

### Drug sensitivity assay

Cells were seeded in 96-well microplates (PerkinElmer); the seeding cell confluency was specifically optimized for each cancer cell line to have cells in growth phase at the end of the assay. After overnight incubation at 37°, cells were treated with DMSO (Merck) for the negative control and with five concentrations of selected drugs in triplicate. Cells were then incubated at 37° for 72hr. Cell viability was assessed by measuring either luminescence with GloMax® Discover instrument from Promega or by nuclei count using the Operetta instrument from PerkinElmer. Luminescence measurements were normalized using background wells as manufacturer protocol. For luminescence measurement, cells were treated with Promega CellTiter-Glo® Luminescent Cell Viability Assay according to the manufacturer protocol. For nuclei count, cells were washed with PBS 1x, fixed with paraformaldehyde (PFA) 4% for 10 min at room temperature, washed with PBS 1x, incubated at room temperature in the dark with HOECHST 33342 (Thermo Fisher Scientific) diluted 1:1000 in PBS 1x for 10 min and finally washed with PBS 1x. Nuclei count was performed using Columbus image analysis software (PerkinElmer). All drug used in this study were purchased from Selleckchem.

## Data availability

Raw sequence data of BC single cell atlas are available on Gene Expression Omnibus (GEO) repository under the accession number.

## Code availability

The code to reproduce main results in the manuscript is available on github at the following address https://github.com/dibbelab/singlecell_bcatlas. Moreover, the single cell atlas can be explored at http://bcatlas.tigem.it.

## Acknowledgments

This work was supported by the STAR (Sostegno Territoriale alle Attività di Ricerca) grant of University of Naples Federico II and the AIRC (Associazione Italiana Ricerca sul Cancro) GRANT MFAG 23162 to GG and by the AIRC (Associazione Italiana Ricerca sul Cancro) Grant IG 2016-18479 to DB and by iPC project H2020 826121 for both GG and DB.

## Author Contribution

GG performed all computational analysis, conceived the method for single-cell drug sensitivity prediction and contributed to the writing of the manuscript. GV implemented the dropseq platform, performed single-cell RNA sequencing and drug response validations. BT performed cytometric analyses, helped with cell culture and RNA-seq library preparation. AI and RB contributed to data discussion and writing of the manuscript. DdB supervised the work, contributed to the writing of the manuscript, and conceived the original idea.

